# Kinetic characteristics of transcriptional bursting in a complex gene model with cyclic promoter structure

**DOI:** 10.1101/2021.12.09.471914

**Authors:** Xiyan Yang, Zihao Wang, Yahao Wu, Tianshou Zhou, Jiajun Zhang

**Affiliations:** School of Financial Mathematics and Statistics, Guangdong University of Finance, Guangzhou 510521, China; Guangdong Province Key Laboratory of Computational Science, Guangzhou 510275, China; School of Mathematics, Sun Yat-Sen University, Guangzhou 510275, China

**Keywords:** Transcription bursting, promoter cycle, multi-ON mechanism, noise, timing

## Abstract

While transcription occurs often in a bursty manner, various possible regulations can lead to complex promoter patterns such as promoter cycles, giving rise to an important issue: How do promoter kinetics shape transcriptional bursting kinetics? Here we introduce and analyze a general model of the promoter cycle consisting of multi-OFF states and multi-ON states, focusing on the effects of multi-ON mechanisms on transcriptional bursting kinetics. The derived analytical results indicate that bust size follows a mixed geometric distribution rather than a single geometric distribution assumed in previous studies, and ON and OFF times obey their own mixed exponential distributions. In addition, we find that the multi-ON mechanism can lead to bimodal burst-size distribution, antagonistic timing of ON and OFF, and diverse burst frequencies, each further contributing to cell-to-cell variability in the mRNA expression level. These results not only reveal essential features of transcriptional bursting kinetics patterns shaped by multi-state mechanisms but also can be used to the inferences of transcriptional bursting kinetics and promoter structure based on experimental data.

## 1. Introduction

Transcription is a core step of gene-expression processes. For most genes in eukaryotic cells, transcription occurs often in a bursty fashion characterized by mRNA synthesis in short periods of activity followed by typically longer periods of inactivity. Single-cell experimental observations have provided evidence for such transcription bursting [1-4]. This variability, which potentially gives rise to cell-to-cell variability in gene expression levels [5-7], can propagate to downstream proteins or target genes through a network. It is believed that the expression noise can not only contribute to alternative cellular fates [8] but also maintain cellular functioning [9-11]. Revealing the mechanism of transcriptional bursting using mathematical models is an important step toward understanding fundamental cellular processes.

Transcriptional bursting kinetics are often characterized by two parameters: burst size and burst timing (or burst frequency). The former is defined as the number of transcripts produced during an active period, whereas the latter involves waiting times between transcription initiation events. For rate-limiting transcription or for transcription with exponential waiting times, mRNA burst sizes follow geometric distributions, which were also observed in early experiments [1, 3, 10, 12]. However, transcription is a complex process involving many regulatory proteins and molecular factors, such as transcription factors, RNA polymerase, and chromatin remodeling complexes [13-16]. Recent experiments have shown that these factors or processes can result in non-exponential waiting-time distribution for genes’ active (ON) and inactive (OFF) durations [17-20]. In addition, some experimental studies have also suggested complex promoter structures [21-23], which often involve nucleosomes competing with or being removed by transcription factors [24], and epigenetic regulation via histone modifications [25-27]. For example, Dunham, *et al*. found that the transcriptional deactivation occurs with graded, stepwise decreases in transcription rate, implying that the gene promoter has variable transcription states [28]. Sepúlveda *et al*. demonstrated that the lysogeny maintenance promoter of phage lambda switches between multiple promoter states [29]. Neuert *et al*. showed that the transcription is a multistate process for the osmotic stress response pathway in *Saccharomyces cerevisiae* [30]. In a word, transcription is finished not in a simple single-step manner but often in a complex multistep fashion [31-33].

It has been verified that mathematical models are a powerful tool for estimating the impacts of molecular mechanisms on transcriptional bursting. In past decades, many theoretical models have been developed to characterize transcriptional bursting kinetics including burst size, burst frequency, and variability in the mRNA abundance [34, 35]. Of these models, the most widely used model is the two-state model [17, 36-39], where the promoter is assumed to switch between transcriptionally active (ON) and inactive (OFF) states with constant switching rates. Queuing models have also been proposed to analyze the impact of transcription regulation on bursting kinetics [31, 40-42], where waiting-time distributions are general (either exponential or non-exponential). For example, Schwabe *et al*. used Erlang-distributed times to model ON-phase and transcription ignition, and found that burst sizes peaked distribution [43]. Kumar *et al*. derived steady-state moments of mRNA distribution and estimated burst parameters for non-geometric burst distribution [44]. Multistate gene models such as multi-OFF models [45, 46], and multi-ON models [47, 48], have been proposed to analyze transcription dynamics. Recently, Zoller *et al*. combined mathematical modeling with experimental measurement to estimate burst parameters in a multi-OFF promoter cycle [49]. In contrast, Rybakova *et al*. used stochastic and ordinary differential equation models to estimate burst parameters in a multi-ON promoter cycle [50]. In addition, Daigle *et al*. inferred the number and configuration of promoter topologies from single-cell gene expression data [51]. In spite of these efforts, how promoter kinetics affect transcription bursting kinetics remain poorly understood. Neither gene models with multi-ON states or with cyclic promoter structure, whose prototypes can be found in natural and synthetic systems [52], have been systematically studied, nor analytical results on burst size and timing have been reported.

In this paper, we introduce and analyze a model of stochastic transcription with cyclic promoter structure (i.e., ON- and OFF states constitute a cycle), focusing on analysis of the effect of multi-ON mechanism on transcriptional bursting kinetics. Interesting results are obtained, e.g., bust size follows a mixed geometric distribution rather than a single geometric distribution that was assumed often in previous studies; each of ON and OFF times obeys a mixed exponential distribution; and the multi-ON mechanism can lead to bimodal burst-size distribution, antagonistic timing of ON and OFF. While our results reveal essential characteristics of transcriptional bursting kinetics, our analysis provides a framework for studying more complex transcription processes.

## 2. Methods

### 2.1 Model hypothesis

Here we introduce a biologically reasonable model for transcription bursting (Fig. 1(a)), which considers the effects of many processes (or factors) such as recruitment of various polymerases and transcription factors, assembly of the pre-initiation complexes, chromatin remodeling, and histone modifications [13-15]. Then, we map this model into a theoretical model of multistate promoter cycle (Fig. (b)) where the promoter proceeds sequentially through several reversible active (ON) states and inactive (OFF) states and returns to the active (ON) state, forming a cycle. We assume that transcription occurs in a bursting manner at each active state but is prohibited at the inactive states (Fig. (c)). Specifically, we assume that the gene promoter has *N* states in total, which consists of *L* active states and *K* inactive states. We call *L* and *K* as active and inactive indexes, respectively. Denoted by *A*_1_, …, *A*_*L*_ ON states and by *I*_1_, …, *I*_*K*_ OFF states. The corresponding biochemical reactions are listed in Table 1.

**Fig. 1.**
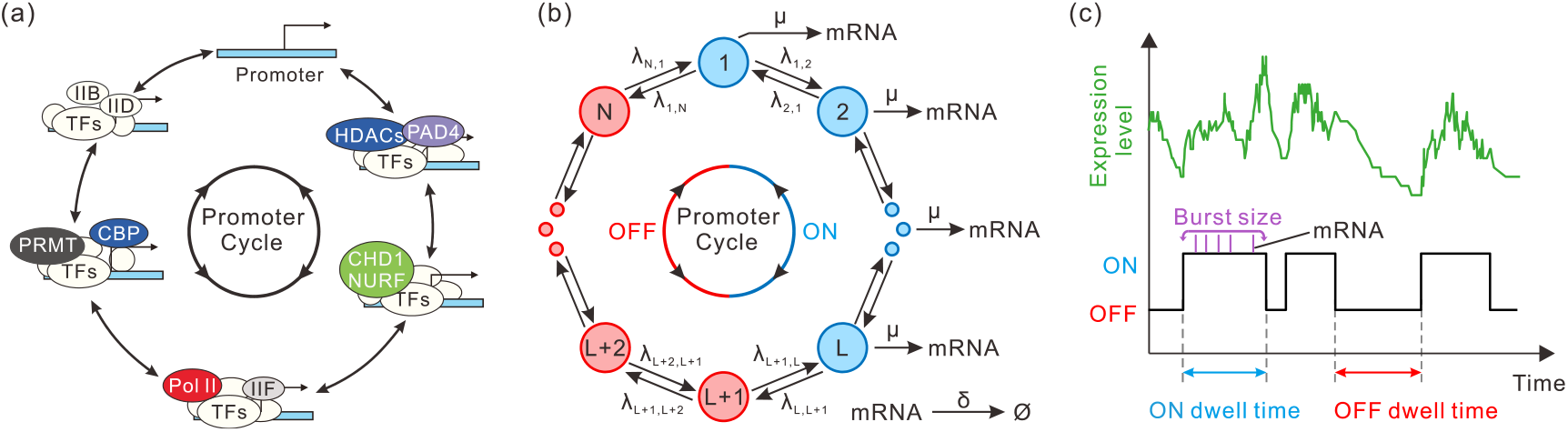
Multistate promoter cycle model for transcriptional bursting. (a) Schematic diagram for a reversible promoter cycle involving recruitments of polymerases and transcription factors, chromatin remodeling, and histone modifications. (b) The reversible promoter cycle in (a) is mapped into a theoretical model, where the blue and red cycles represent transcription active (ON) and inactive (OFF) states, respectively. (c) Representative time series for mRNA expression and gene activity, where the gene transcribes in the ON state in a bursting manner.

**Table 1.**
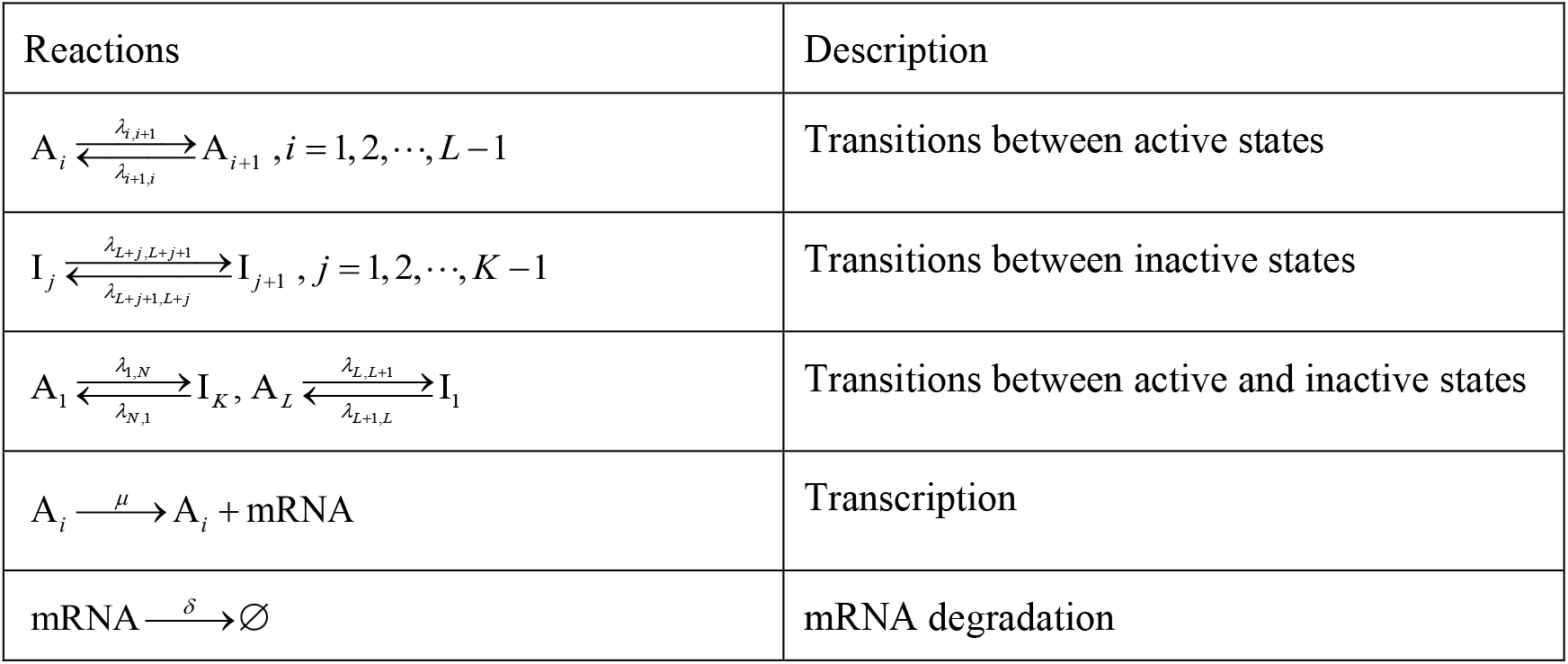
Biochemical reactions for the transcription model to be studied.

| Reactions | Description |
| --- | --- |
| $A_i \xrightleftharpoons[\lambda_{i+1,i}]{\lambda_{i,i+1}} A_{i+1}, i = 1, 2, \dots, L-1$ | Transitions between active states |
| $I_j \xrightleftharpoons[\lambda_{L+j+1,L+j}]{\lambda_{L+j,L+j+1}} I_{j+1}, j = 1, 2, \dots, K-1$ | Transitions between inactive states |
| $A_1 \xrightleftharpoons[\lambda_{N,1}]{\lambda_{1,N}} I_K, A_L \xrightleftharpoons[\lambda_{L+1,L}]{\lambda_{L,L+1}} I_1$ | Transitions between active and inactive states |
| $A_i \xrightarrow{\mu} A_i + \text{mRNA}$ | Transcription |
| $\text{mRNA} \xrightarrow{\delta} \emptyset$ | mRNA degradation |

Note that both *L* = 1 and *N* > 2 correspond to the multi-OFF model (i.e., the promoter has more than one OFF state but only one ON state), and both *K* = 1 and *N* > 2 correspond to the multi-ON model (i.e., the promoter has more than one ON state but only one OFF state). Therefore, the model introduced here includes transcription models that were previously studied [45-48], and can be used to model general transcription dynamics generated by gene promoters with multiple states. We point out that for the multi-OFF and multi-ON models, some analytical results on steady-state mRNA distribution and noise have been obtained [45, 47], but for multistate promoter models, the analytical expressions on transcription bursting kinetics, such as distributions of burst size and timing, have not been derived so far.

### 2.2. Analysis framework

#### 2.2.1. Burst size distribution

Let *U* represent the promoter’s active state, which is a time-dependent random variable, i.e., *U* (*t*) = *u* ∈{*A*_1_,, *A*_*L*_}, *B* (*t*) be burst size, i.e., *B* (*t*) = *n, n* = 0,1, 2,…, and *T* represent the exit time (i.e., burst termination time) from an active state. We assume that at time *t* = 0, the promoter begins to switch to active state *A*_1_ from inactive state *I*_*K*_, or to active state *A*_*L*_ from inactive state *I*_1_. Let *S*_*n,u*_ (*t*) represents the survival probability that *n* mRNAs are produced per burst in state *u* at time *t*. Then, the transcriptional system’s state at time *t* can be described as *S*_*n,u*_ (*t*) = Pr ob.{*T* > *t X* = *n,U* (*t*) = *u*}, where we stipulate *S*_*n,u*_ (*t*) ≡ 0(*n* < 0). Note that the probability that burst termination time *T* falls within an infinitesimal interval (*t, t* + Δ*t*) can be expressed as

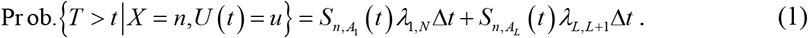

Therefore, the joint probability density function *P*_burst_ (*n,t*) for burst size *X* (a discrete random variable) and burst termination time *T* (a continuous random variable) includes all incoming fluxes driving the system from active state *A*_1_ to inactive state *I*_*K*_, or from active state *A*_*L*_ to inactive state *I*_1_, that is, 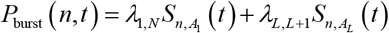. Then, the steady-state probability distribution for burst size *X* is calculated according to

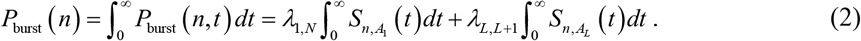

The key is how 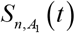 and 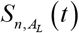 are obtained. However, if we rewrite 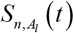 as *S*_*n,l*_ (*t*) for convenience, the master equation for survival probability *S*_*n,l*_ (*t*) can be described as

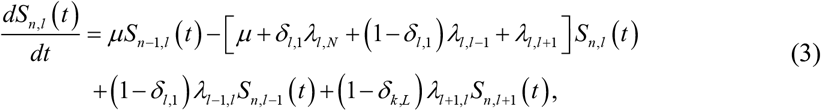

Where *δ*_*i, j*_ is a Kronecker delta. Note that this equation can be solved by introducing probability-generating functions. Specifically, if defining 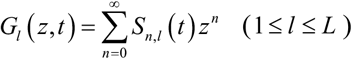 and denoting **G** (*z, t*) = (*G*_*1*_ (*z, t*),, *G*_*L*_ (*z, t*))^*T*^, we can derive a linear partial differential equation group of the form: *∂***G** (*z, t*) *∂t* = **H** (*z*)**G** (*z, t*), where coefficient matrix **H**(*z*) is a tridiagonal matrix with the diagonal elements taking the form *h*_*l*_ (*z*) = *µ* (*z* −1) − *δ*_*l*, 1_*λ*_*l, N*_ − (1− *δ*_*l*, 1_)*λ*_*l*, *l* −1_ − *λ*_*l*, *l* +1_ (1 ≤ *l* ≤ *L*) (seeing Appendix A for details).

#### 2.2.2. Dwell time distributions and burst frequency

Note that the distribution of the time that the promoter dwells on active (or inactive) state, denoted by *f* ^ON^ (*t*) (or *f* ^OFF^ (*t*)), is equal to the sum of the times dwelling on all ON (or OFF) states. In order to calculate *f* ^ON^ (*t*) and *f* ^OFF^ (*t*), we introduce two column vectors 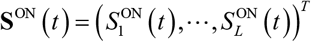 and 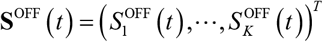, where 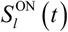 and 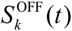 are the survival probabilities that the promoter is still at respectively. *l*^*th*^ ON state and *k*^*th*^ OFF state, respectively.

Now, we establish the master equation for 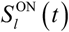 and 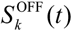. Denote by **T**^ON^ (*L* × *L* matrix) and **T**^OFF^ (*K* × *K* matrix) the transition matrices of promoter among the active states and among the inactive states respectively, and introduce two matrices 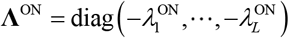 and 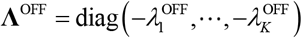, where 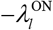 and 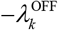 are eigenvalues of matrices **T**^ON^ and **T**^OFF^, respectively. Then, the master equations of the survival probability for ON and OFF states can take the following unified form

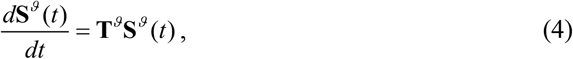

Where *ϑ* = ON or OFF.

Eq. (4) is a linear equation group and is easily solved. In order to derive the analytical expression of **S**^*ϑ*^ (*t*), we introduce two diagonal matrices 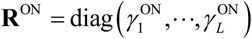 where 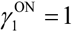 and 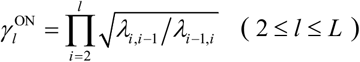, and 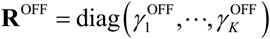 where 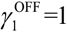 and 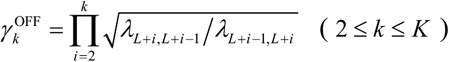. In addition, we define 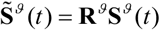 and 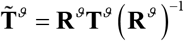, where *ϑ* = ON or OFF. Note that there is an orthogonal matrix 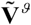 such that 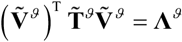. Then, vector **S**^*ϑ*^ (*t*) for the survival probabilities can be analytically expressed as

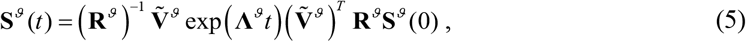

Where **S**^*ϑ*^ (0) is the vector of initial survival probabilities determined by the steady-state distribution of the promoter states. To that end, the total dwell time distribution at the active and inactive states can be calculated according to

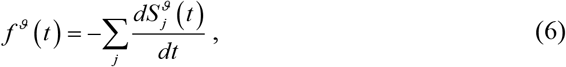

where *ϑ* = ON or OFF.

After having obtained two dwell time distributions at ON and OFF states, we can separately calculate the mean ON and OFF times, denoted by ⟨*τ* ^ON^ ⟩ and ⟨*τ* ^OFF^ ⟩. Furthermore, the mean burst frequency (BF) can be formally expressed as

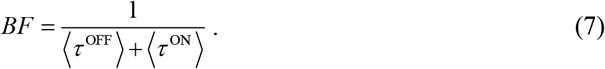

#### 2.2.3. mRNA mean and noise

Note that the master equation for mRNA is different from that for burst size. Here we first establish the master equation for the former and then calculate the mean and noise of mRNA.

Let *P*_*s*_ (*m*;*t*) (1 ≤ *s* ≤ *N*) represent the probability that the number of mRNA molecules is at state *A*, …, *A, I*, …, *I* at time *t*, respectively. Set **P** (*m*;*t*) = [*P*_*1*_ (*m*;*t*),…, *P*_*N*_ (*m*;*t*)]^T^, and denote by **A =** (*λ*_*k,l*_) the *N* × *N* transition matrix of a promoter. Note that *λ*_*k,l*_ = 0 means that there is no transition occurrence from state *k* to state *l*. Let **Ω** = diag(*µ, µ*,…, *µ*, 0, 0,…, 0) represent the transcription matrix. Then, the corresponding chemical master equation describing the mRNA dynamics takes the following form:

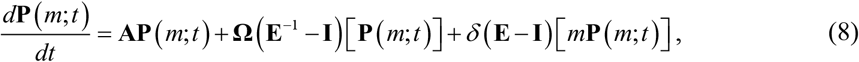

where **E, E**^−1^ are shift operators and **I** is the identity operator.

Eq. (8) can be solved using the binomial moment (BM) method that we previously developed [53]. The framework of this method is stated below. If the factorial BM for the probability *P*_*s*_ (*m*;*t*) is defined as 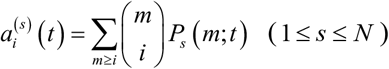, then the total BM for the total probability defined as 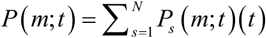 is calculated by 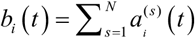. Note that *b*_0_ (*t*) = 1 due to the conservative condition: 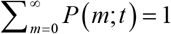. Furthermore, if we introduce column vector 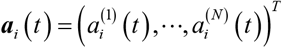, it follows from Eq. (8) that

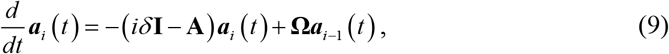

which is a linear ordinary differential equation group and is easily solved. Once all 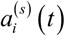 and further the total BMs *b*_*i*_ (*t*) are obtained, the mRNA probability distribution can be reconstructed according to

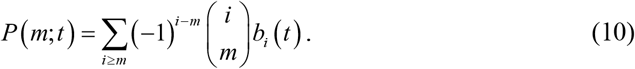

In addition, if the mRNA noise is quantified by the ratio of the variance over the squared mean (called noise intensity and denoted by 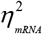, then 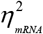 is calculated according to

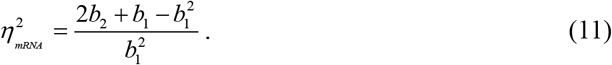

## 3. Analytical results

Here we present analytical results, obtained according to the above analysis framework, to help understand the mechanism of how the cyclic promoter structure affects transcriptional bursting kinetics.

### 3.1. General results

First, we present analytical results for burst size distribution. If we let 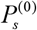 represent the steady-state probability that the promoter is in state *s*, where *s* = 1, 2,…, *N*, then these probabilities can be analytically derived (seeing Appendix B for details). Interestingly, we find that initial conditions *S*_*n,l*_ (0) are directly related to the steady-state distributions at the OFF states. Specifically, 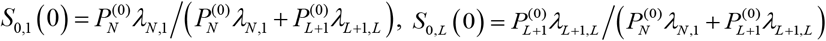, and *S*_*n,l*_ (0) = 0 for other subscripts. Owing to the special structure of matrix **H**(*z*), generating function *G*_*l*_ (*z, t*) can be analytically expressed as

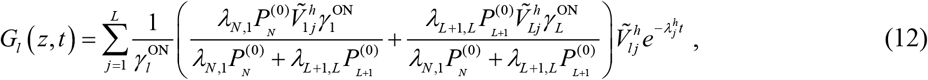

Where 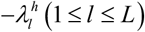 are eigenvalues of matrix **H** (*z*) and are thus functions of *z*, and 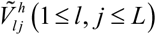 are the elements of orthogonal eigenvector 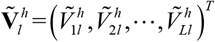 associated with the eigenvalues 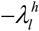. Furthermore, generating function *G*_burst_ (*z*) for burst size distribution *P*_burst_ (*n*) can be expressed as

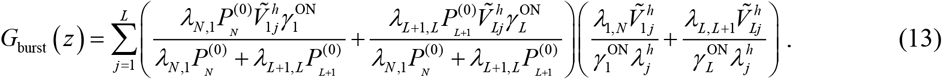

Thus, burst size distribution *P*_burst_ (*n*) can be given via 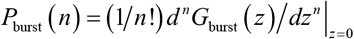.

Second, we present analytical results for dwell time distributions. Note that 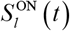 and 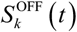 for survival probabilities at the *l*^*th*^ active state and at the *k* ^*th*^ inactive state can be respectively expressed as

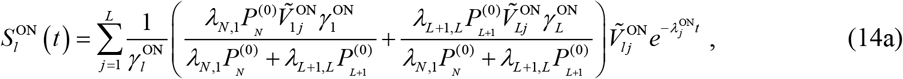

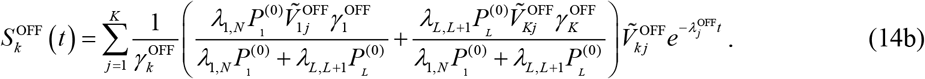

where 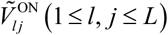 and 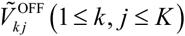 are the elements of matrices 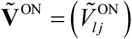 and 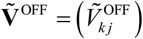, respectively.

Using Eq. (14a) and (14b), the dwell time distributions are thus expressed as

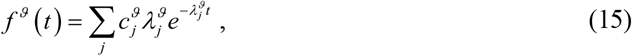

where if *ϑ* = ON, then

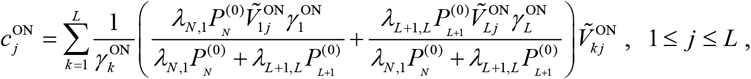

and if *ϑ* = OFF, then

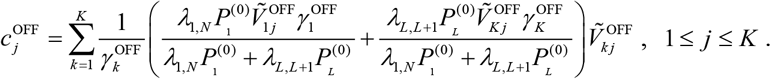

Eq. (15) indicates that the dwell time distribution at ON or OFF states are in general the sum of exponential distributions.

Third, we present analytical results for BMs. We find that the steady-state total BMs take the following form

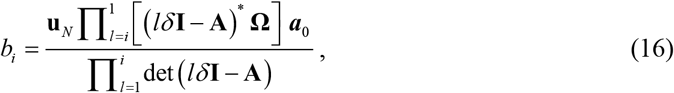

Where **u**_*N*_ = (1,1,…,1) is a *N* − dimensional row vector, (*iδ* **I** − **A**)^*^ and det (*iδ* **I** − **A**) are respectively the adjacency matrix and the determinant of matrix (*iδ* **I** − **A**), and the initial vector ***a***_0_ is just the steady-state distribution of promoter states (see Appendix B).

In particular, the first-order binominal moment, i.e., the mRNA mean is given by (see Appendix C)

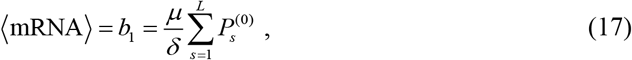

and the second-order binomial moment takes the form

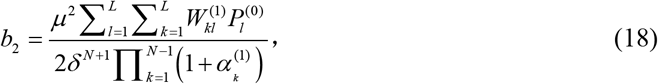

Where 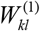 represents a determinant of an (*N* −1)×(*N* −1) matrix that is the left part of matrix (*δ* **I** − **A**) by crossing out its *k*^*th*^ row and *l*^*th*^ column elements, and 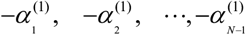 are the nonzero eigenvalues of matrix **A** *δ*. Then, we can show that the Fano factor for mRNA, defined as the ratio of the variance over the mean, is in general not equal to one, implying that the mRNA distribution is in general not Poissonian but may be super-Poissonian or sub-Poissonian.

We point out that all the above analytical results are formal and tell us the finite information on the effect of the cyclic promoter structure on transcriptional bursting kinetics. In order to clearly see how the cyclic promoter structure affects transcriptional bursting kinetics, we will consider special cases.

### 3.2. Specific results

Here we consider two representative cases: (1) Multi-OFF mechanism [45], i.e., *L* = 1 and *K* = *N* −1 ; (2) Mixture mechanism where all the forward (i.e., clockwise) transition rates of active states are identical (denoted by *λ*_*f*_) and all the backward (i.e., counterclockwise) transitions of active states are also identical (denoted by *λ*_*b*_); and all the forward transition rates of inactive states are the same (denoted by *ρ* _*f*_ *λ*_*f*_ with *ρ* _*f*_ being a common forward ratio) whereas all the backward transition rates of inactive states are also the same (denoted by *ρ*_*b*_ *λ*_*b*_ with *ρ*_*b*_ being a common backward ratio).

For the multi-OFF mechanism, according to the above general formula, it is not difficult to show that the generating function for burst size is given by

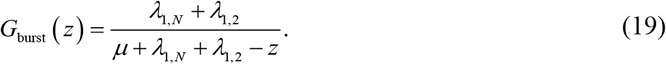

Thus, the burst size distribution takes the form

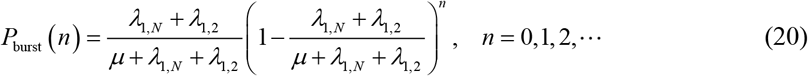

which is a geometric distribution. These results are in agreement with previous assumptions [43].

For the mixture mechanism, we denote 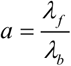 and 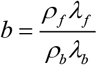. Using the above formulae and noting the fact: 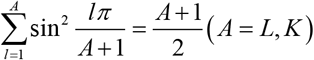, we can show

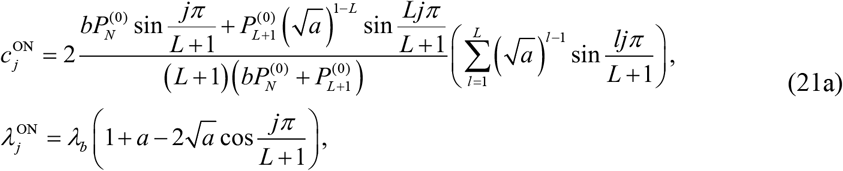

where *j* = 1, 2,…, *L*, and

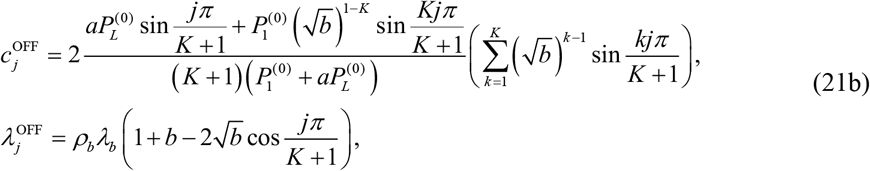

where *j* = 1, 2,…, *K*. Correspondingly, generating function *G*_burst_ (*z*) for burst size is given by (see Appendix D for derivation)

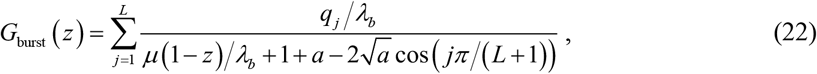

where

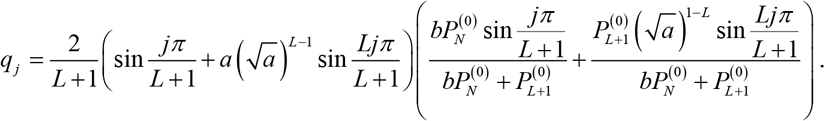

Thus, the bust size distribution takes the form

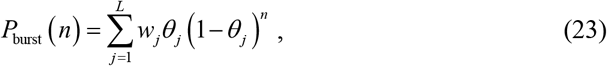

where *n* = 0,1, 2, ,

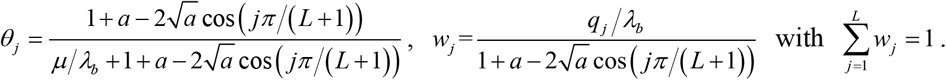

Eq. (23) indicates that burst size distribution is a mixture of geometric distributions. At the same time, the mean burst size is given by

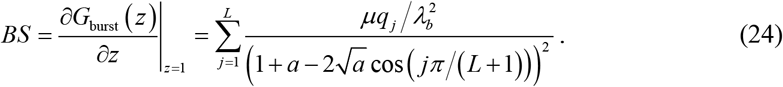

In addition, we can also calculate the burst size noise, which is given by

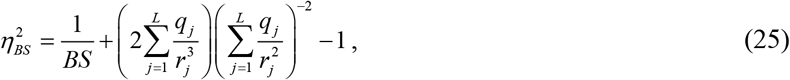

where 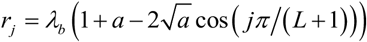. The corresponding noise will contribute to the mRNA noise.

## 4. Numerical Results

In contrast to previous studies that focused on the effect of multi-OFF mechanisms on gene expression, here we numerically demonstrate influences of multi-ON mechanisms on transcriptional bursting kinetics.

### 4.1. Multi-ON mechanism can lead to bimodal burst size distribution

Busting kinetics is commonly characterized by burst size (BS) and burst frequency (BF). Here, we analyze how the multi-ON mechanism affects burst size and its distribution. Previous studies of the two-state gene model assumed that burst size follows a geometrical distribution [1, 12], but whether or not this assumption is reasonable is unclear. In particular, it is not clear whether the burst size still obeys a geometric distribution in the case of multi-ON mechanisms. By performing numerical calculations, we find that multi-ON mechanism can induce bimodal distribution of burst sizes, referring to Fig. 2(a)-(c). Fig. 2(a) and (b) show two representative unimodal and bimodal burst size distributions, respectively, and Fig. 2(c) shows how the number of the most probable mRNA molecules obtained by a statistical method depends on the active index (*L*). In Fig. 2(a) and (b), histograms are stochastic simulations using the Gillespie stochastic simulation algorithm (SSA) [54], and solid lines are theoretical predictions. Fig. 2(c) shows that the bimodal mRNA distribution exists for a large range of *L*. Specifically, the burst size distribution is unimodal at the origin with *L* = 1 (i.e., multi-OFF model), a geometric distribution as predicted by Eq. (20). Then, the burst size distribution becomes a single peak away from the origin if we increase *L*, e.g., *L* = 2, 3. Finally, the burst size distribution becomes bimodal with one peak at the origin and another peak away from the origin for larger *L* with *L* ≥ 4. These results imply that burst size distribution in the case of the multi-ON mechanism is non-geometric but a mixture of geometric distributions as predicted by Eq. (23).

**Fig. 2.**
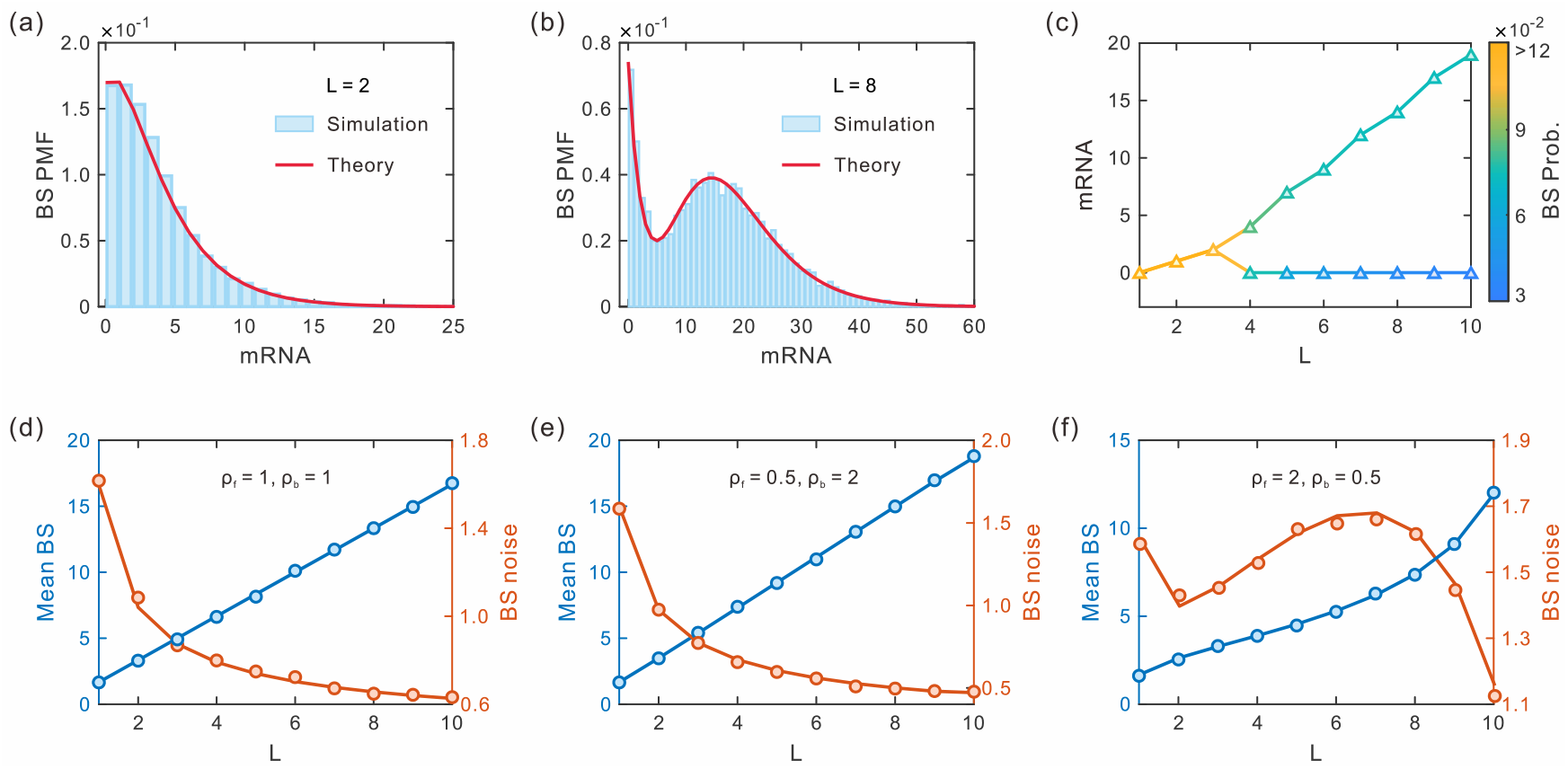
Influence of multi-ON mechanism on burst size. (a) and (b) Two representative burst-size distributions corresponding to *L* = 2 and *L* = 8 ; (c) Dependence of the most probable mRNA numbers on active index *L*, where the color bar represents the BS probability at the most probable mRNA numbers. In (a)-(c), *ρ* _*f*_ = 0.5, *ρ*_*b*_ = 2. (d), (e) and (f) Dependence of mean BS (red) and BS noise (blue) on *L* for three representative cycles with *ρ* _*f*_ = 1, *ρ*_*b*_ = 1, *ρ* _*f*_ = 0.5, *ρ*_*b*_ = 2, and *ρ* _*f*_ = 2, *ρ*_*b*_ = 0.5, where the solid lines represent theoretical results and circles are numerical results obtained by SSA [54]. Parameter values are set as *λ*_*f*_ = 0.1, *λ*_*b*_ = 0.5, *µ* = 1,*δ* = 0.1, *N* = 11.

Next, we analyze the influences of the multi-ON mechanism on mean burst size and burst size noise, referring to Fig. 2(d)-(f), where three representative promoter loops are shown. In all loops, we observe that mean burst size is always a monotonically increasing function of the active index (*L*), implying that more active states can lead to a larger burst size. In addition, we observe that burst size noise is a monotonically decreasing function of the active index in the loops where the forward ratio is equal to the backward ratio, and the forward ratio is smaller than the backward ratio. However, there is a non-monotonic relationship between the active index and burst size noise if the forward ratio is larger than the backward ratio. These results indicate that burst size noise in the case of the multi-ON mechanism can exhibit different characteristics.

Previous studies showed that genes with a multistep mechanism could have peaked burst size distribution and reduce noise in burst size [43]. Here, we point out that the multi-ON mechanism can induce more complex behaviors of bust size, such as bimodal bust size distribution and non-monotonic burst size noise.

### 4.2. Multi-ON mechanism can lead to antagonistic timing of ON and OFF

Here we focus on analyzing how the multi-ON mechanism influences burst timing between transcription initial events by performing numerical calculations, referring to Fig. 3. The first and second rows of Fig. 3 show two representative OFF dwell-time distributions (Fig. 3(a) and (b)) and ON dwell-time distributions (Fig. 3(d) and (e)), as well as how the times for peak appearance in OFF dwell-time distribution (Fig. 3(c)) and ON dwell-time distribution (Fig. 3(f)) depend on the active index (*L*), respectively. First, we observe that OFF dwell-time distribution changes from unimodality to bimodality, whereas ON dwell time distribution varies from bimodality to unimodality with the increase of *L* (Comparing Fig. 3(a)-(c) with Fig. 3(d)-(f)). For example, the OFF dwell time distribution is bimodal with one peak at the origin and another peak away from the origin for a broad range of *L* with 1 ≤ *L* ≤ 8, and then this distribution becomes unimodal with a single peak away from the origin if *L* = 9, and further becomes unimodal with a peak at the origin if *L* = 10 (see Fig. 3(c)). However, for the ON dwell time, the distribution is unimodal with a peak at the origin if *L* = 1, and is an exponential distribution corresponding to a multi-OFF model. And the distribution changes from unimodality with one single peak away from the origin if *L* = 2 to bimodality with one peak at the origin and another peak away from the origin for a broader range of *L* : 3 ≤ *L* ≤ 10 (see Fig. 3(f)).

**Fig. 3.**
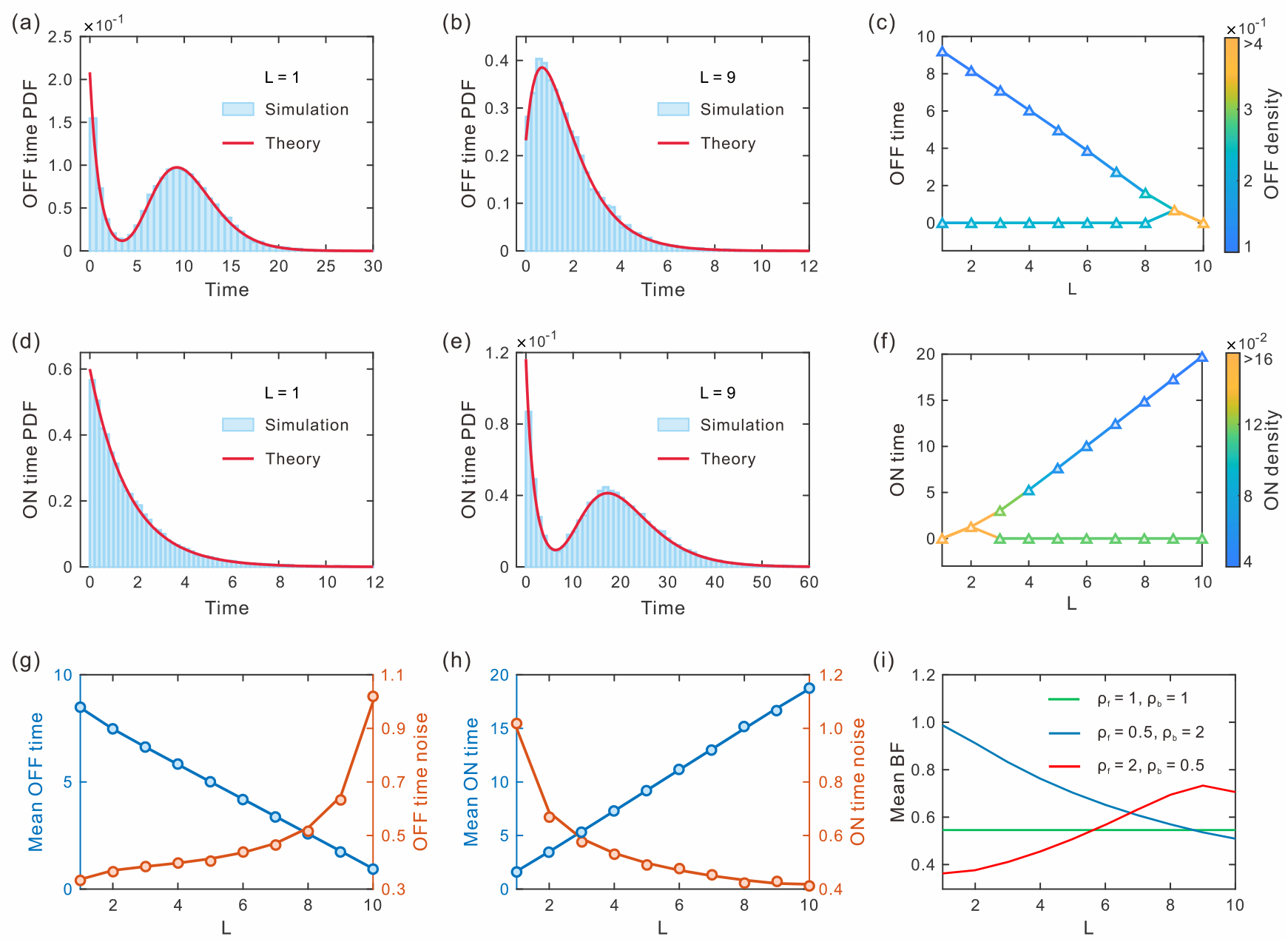
Influence of multi-ON mechanism on burst timing. (a) and (b) Two representative OFF dwell-time distributions corresponding to *L* = 1 and *L* = 9 ; (c) Dependence of the time for peak appearance in OFF dwell time distributions on the active index (*L*); (d) and (e) Two representative ON dwell-time distributions for *L* = 1 and *L* = 9 ; (f) Dependence of the time for peak appearance in ON dwell time distributions on *L*. In (a)-(f), *ρ* _*f*_ = 0.5, *ρ*_*b*_ = 2. (g) Dependence of the mean OFF time and OFF time noise on *L* ; (h) Dependence of mean ON time and ON time noise on *L* ; (i) Dependence of mean BF on *L* for three representative loops : *ρ* _*f*_ = 1, *ρ*_*b*_ = 1 (green line), *ρ* _*f*_ = 0.5, *ρ*_*b*_ = 2 (blue line), and *ρ* _*f*_ = 2, *ρ*_*b*_ = 0.5 (read line). In (g) and (h), the solid lines represent theoretical results and circles are numerical results obtained by SSA [54]. Parameter values are set as *λ*_*f*_ = 0.1, *λb* = 0.5, *µ* = 1,*δ* = 0.1, *N* = 11.

The bottom row shows the mean and noise of OFF dwell time (Fig. 3(g)) and ON dwell-time (Fig. 3(h)), respectively, as well as mean burst frequency for three representative loops (Fig. 3(i)). We observe that with the rise of the active index, the mean OFF time decreases whereas mean ON time increases, but the OFF time noise increases and ON time noise decreases. In addition, we find that with increasing the active index, the mean burst frequency can exhibit different characteristics, such as no change with *ρ* _*f*_ = *ρ*_*b*_ = 1, decrease with *ρ* _*f*_ = 0.5, *ρ*_*b*_ = 2, and increase with *ρ* _*f*_ = 2, *ρ*_*b*_ = 0.5.

The above analysis implies that the multi-ON mechanism can give rise to antagonistic timing of ON and OFF, and induce different characteristics of burst frequency, hinting that the multi-ON mechanism has an un-negligible influence on transcriptional bursting timing.

### 4.3. Propagation of bursting kinetics to mRNA variability

Single-molecule experiments have verified that mRNAs are produced often in a bursting manner [1, 2]. It is of interest to understand how bursting kinetics affect the variations in mRNA levels. Previous studies focused on the influence of burst parameters such as mean burst size and burst frequency on gene expression levels and noise by analyzing queuing models [31, 32, 43, 44]. However, how the burst size and timing affect gene expression dynamics in a multistate promoter remains elusive, referring to Fig. 4(a) and (b). Here, we explore the influences of burst size distribution and dwell-time distribution on mRNA distribution, and the numerical results are shown in Fig. 4(c)-(j).

**Fig. 4.**
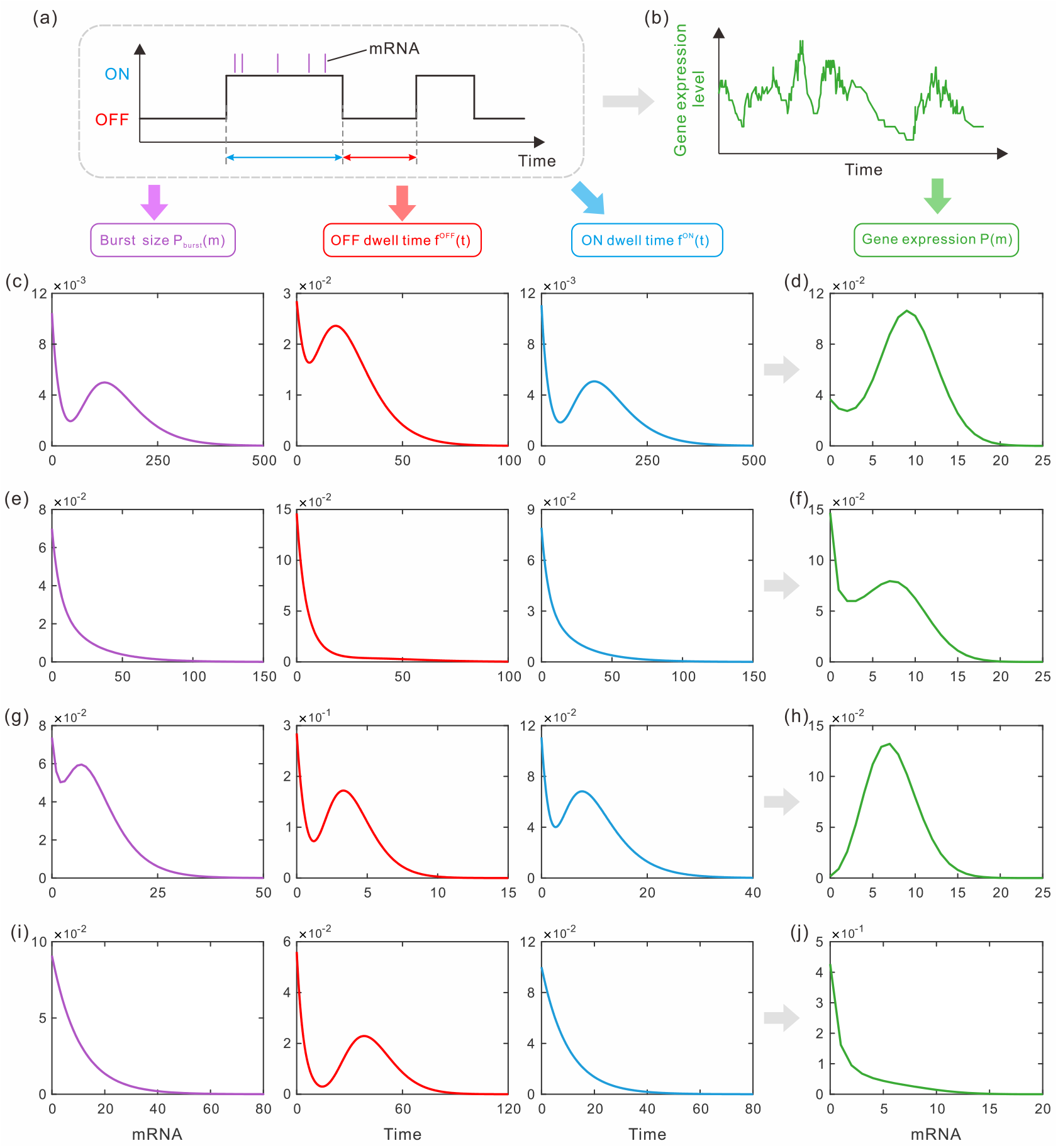
Influence of transcriptional bursting kinetics on gene expression dynamics. (a) Schematic of a multistate promoter model of transcriptional bursting, where the promoter switches between multi-ON states and multi-OFF states, and transcribes only in the ON state; (b) Time series change in the number of mRNA counts. (c), (e), (g) and (i) Burst size distribution (the first column), OFF dwell-time distribution (the second column), ON dwell-time distribution (the third column) for different parameter values; (d), (f), (h) and (j) mRNA distribution (the last column) for different parameters. Parameter values are set as:*µ* = 1,*δ* = 0.1. *ρ* _*f*_ = 0.5, *ρ*_*b*_ = 3, *N* = 11. Other parameters are: (c, d) *λ*_*f*_ = 0.01, *λb* = 0.05, *L* = 7; (e, f) *λ*_*f*_ = 0.08, *λb* = 0.06, *L* = 3; (g, h) *λ*_*f*_ = 0.1, *λb* = 0.5, *L* = 5;(i, j) *λ*_*f*_ = 0.02, *λb* = 0.08, *L* = 1.

From Fig. 4(c) and (d), we first observe that the burst size distribution, OFF dwell-time distribution, and ON dwell-time distribution are bimodal, and the mRNA distribution is also bimodal. From Fig. 4(e) and (f), we find that the burst size distribution, OFF dwell-time distribution, and ON dwell-time distribution are unimodal, but the mRNA distribution is bimodal. These results indicate that both unimodal and bimodal distributions of burst size and dwell-time can lead to bimodal mRNA distribution. On the other hand, we find that the bimodal or unimodal distributions of burst size, OFF dwell-time, and ON dwell-time can lead to unimodal mRNA distribution, referring to Fig. 4(g)-(j). A previous study showed that unimodal or bimodal gene expression has important biological implications, i.e., a cause of phenotypic diversity in a population of genetically identical cells [55]. These results indicate that the variation in the distributions of burst size and dwell-time can give rise to variability in the distribution of gene expression production and further phenotypic variation.

Next, we analyze the influence of burst size noise, OFF dwell-time noise, and ON dwell time noise on mRNA noise, with numerical results shown in Fig. 5. We observe that the changes in burst size noise, OFF dwell time noise, and ON dwell time noise have different characteristics such as increasing, decreasing, and non-monotonic with the increase of the active index (*L*), referring to Fig. 5(a), (c), and (e). However, the mRNA noise is always a monotonically decreasing function of *L* (see the last column of Fig. 5). These results indicate that the integration of the burst size noise and dwell-time noise may reduce gene expression noise. Our previous study also showed this by analyzing multi-OFF models [45]. Together, the results imply that a multistep process is a universal mechanism of reducing gene expression noise.

**Fig. 5.**
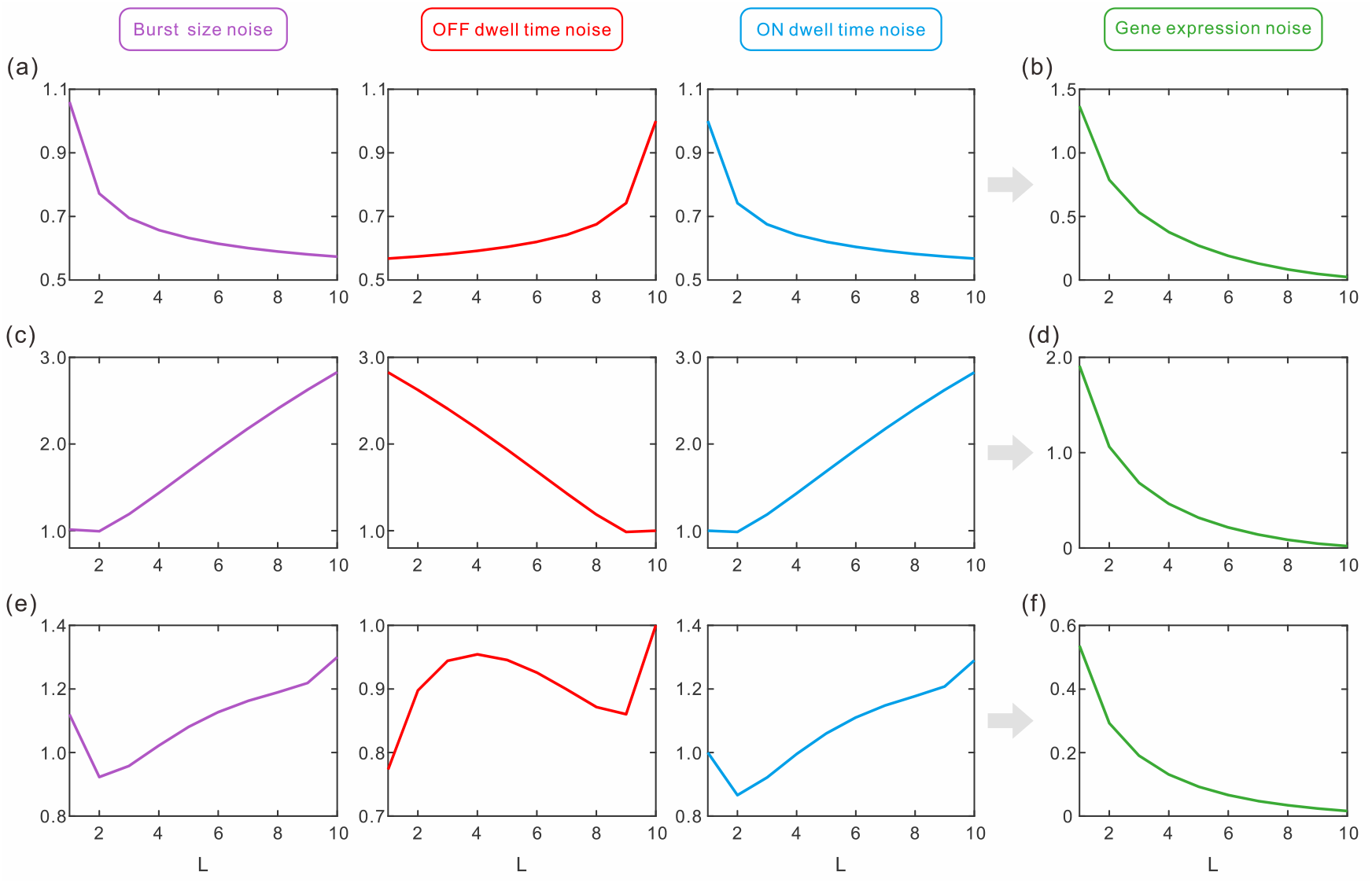
Influence of multi-ON mechanism on four kinds of noise for different parameter values. (a), (c) and (e) Burst size noise (the first column), OFF dwell time noise (the second column), ON dwell time noise (the third column); (b), (d), and (f) Gene expression noise. Parameter values are set as: (a, b) *λ*_*f*_ = 0.1, *λb* = 0.5, *ρ* _*f*_ = 1, *ρ*_*b*_ = 1; (c, d) *λ*_*f*_ = 0.08, *λb* = 0.06, *ρ* _*f*_ = 2.5, *ρ*_*b*_ = 2.5; (e, f) *λ*_*f*_ = 0.4, *λb* = 0.8, *ρ* _*f*_ = 0.5, *ρ*_*b*_ = 1.5. Other parameter values are *µ* = 10, *δ* = 0.1, *N* = 11.

The above analysis shows that the multi-ON mechanism can induce complex transcription kinetics, e.g., unimodal or bimodal distribution of mRNA counts, different burst sizes and timing, and different characteristics of noise. The obtained results imply that the propagation of bursting kinetics to gene expression is complex, and would be important for understanding cellular processes and variation in cell populations.

## 5. Discussion and conclusion

Transcription is an inherently stochastic process and often occurs in a bursting fashion. Because of involving recruitment of TFs and DNA polymerases, chromatin remodeling, and sequence of transitions between activity states of the promoter, the kinetics of transcriptional bursting is poorly understood. Previous studies showed that mRNA burst size distribution is geometric with exponential waiting time for two-state promoter models [37, 39], but how transcription burst size and timing in multistate promoter models remains unclear.

In the present work, we have analyzed a gene model that corporate the complexity of promoter structure, focusing on the effects of multi-ON mechanisms on transcriptional bursting kinetics. By establishing the survival probability master equations for mRNA bursting production and promoter dwelling at ON/OFF states, we have successfully derived analytical expressions for burst size distribution and ON/OFF waiting time distributions, and have found that the burst size is a mixed geometric distribution and the ON/OFF dwell time is a mixed exponential distribution in the case of the mixture mechanism. These obtained distributions can be used to analyze transcriptional burst size and timing for arbitrarily complex promoter structure shown in Fig. 1.

While previous studies on multi-OFF models [46] and multi-ON models [47] focused on analysis of mRNA distribution and transcriptional noise, in the present work we have explored the effects of the multi-ON mechanism on transcriptional bursting kinetics. We have shown that this mechanism can lead to bimodal burst size distribution and different patterns of burst size noise, which is different from previous results that the multistep mechanism can give rise to a peaked burst-size distribution and a reduction in the noise of burst size [43]. In addition, we found that the multi-ON mechanism can lead to antagonistic timing of ON and OFF. For example, the OFF waiting time distribution changes from bimodality to unimodality, whereas the ON waiting time distribution changes from unimodality to bimodality, as the active index increases. Finally, by exploring the distributions and noise for bust size, OFF/ON dwell time, and mRNA copy-number, we found that the propagation of bursting kinetics to mRNA variability is complex. In contrast to previous studies on two-state gene models, which showed that the lower expression noise could result from higher burst frequency, or smaller burst size [34, 39, 56], here we showed that the qualitative conclusions no longer hold in multistate promoter models. More specifically, we find that mRNA noise always reduces, but the bust size noise, OFF/ON dwell time noise, and burst frequency can exhibit complex behaviors in the case of the multi-ON mechanism.

Recently, many studies aimed at the inference of kinetic parameters from experimental data on single cells [49, 51]. In a previous study, we showed that different ON/OFF waiting time distributions can exhibit identical steady-state mRNA distributions [47], implying that when one infers a gene system based on transcription dynamics obtained by measurements of fluctuations of mRNA levels, caution must be kept. However, our results obtained here would be used to infer the promoter structure and investigate the variability of the transcription output based on the experimental data. It should be pointed out that the intermediate processes of gene expression such as the partitioning of mRNA at cell division [57], the alternative splicing [58], and the feedback loop [59], would additionally influence the obtained results, but they are neglected in our study. How details of these factors affect transcriptional bursting kinetics is worth further investigation.

## Acknowledgments

This work was supported by the National Nature Science Foundation of P. R. China (Nos. 11601094, 12171494, 11631005, 11775314, 11931019), Guangdong Key Research and Development Project (2019B0233002), and Guangdong Province Key Laboratory of Computational Science at the Sun Yat-sen University (2020B1212060032).

## Conflict of interests

The authors have declared that no competing interest exists.

## Appendix

### A. Expression for coefficient matrix **H**(z)

According to the definition of generating function and Eq. (3) in the main text, we can show the expression for coefficient matrix

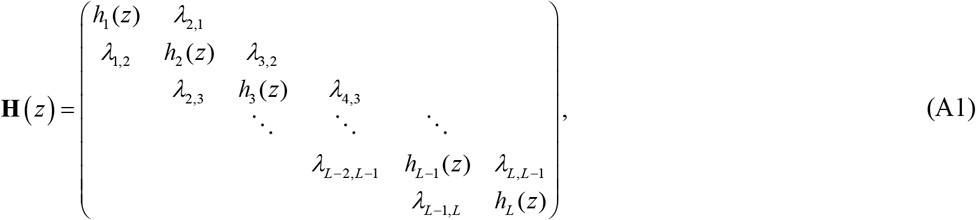

Where *h*_*l*_ (*z*) = *µ* (*z* −1) − *δ*_*l*1_*λ*_*l,N*_ − (1− *δ*_*l*1_)*λ*_*l,l*−1_ − *λ*_*l,l*+1_, *l* = 1, 2,…, *L*.

### B. Derivation of steady-state probability for promoter state

According to Table 1 in the main text, we can establish the following master equation for promoter kinetics

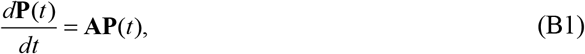

Where **P**(*t*) = (*P*_1_ (*t*), *P*_2_ (*t*),, *P*_*N*_ (*t*))^T^ with *P* (*t*)_*k*_ being the time-dependent probability of promoter at state *k*, and **A** is the transcription matrix of promoter state. Our interest is in deriving the steady-state probability 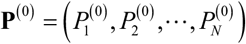 satisfying the condition: **AP**^(0)^ = 0.

First, note that the transition matrix **A** is an M-matrix (i.e., the sum of every column elements is equal to zero). So we can assume the eigenvalues of the matrix **A** as 0, −*α*_1_, −*α*_2_ …, −*α*_*N* −1_ with *α*_*s*_ ≠ 0, and the eigenvalues of the matrix **A**_*k*_ (i.e., crossing out the *k*^*th*^ row and *k*^*th*^ column entry of matrix **A**) as 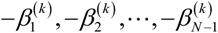 with 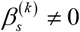. This assumption implies that the characteristic polynomial of **A** and **A**_*k*_ takes the following form, respectively

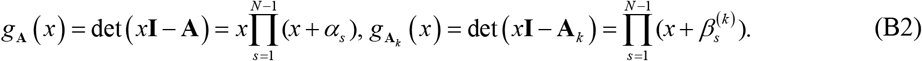

Note that Laplace’s formula for the M-matrix **A** gives

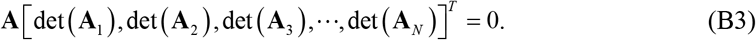

Since the null space of **A** is one-dimensional, we have **P**^(0)^ = *ρ* [det (**A**_**1**_), det (**A**_**2**_),…, det (**A**_***N***_)]^T^. From the conservative condition **u**_***N***_ **P**^(0)^ = 1 with **u**_*N*_ = (1,1,…,1) being a row vector, we can obtain

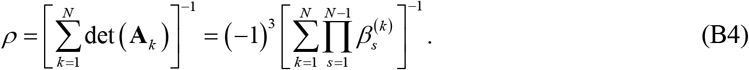

From Eq. (B2), we have 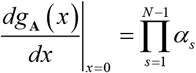. On the other hand, Jacobi’s formula gives

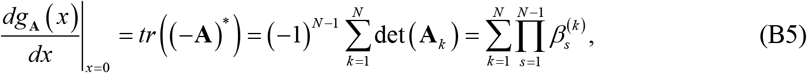

where “tr” represents the trace of a matrix. The combination of both yields the equality 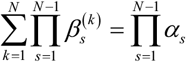. Finally, we obtain the explicit expression of steady-state probability for promoter at every state

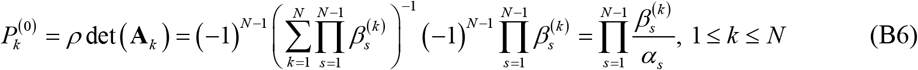

### C. Derivation of mRNA mean and noise

First, we compute the binomial moments (BMs). Let 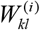 represent a determinant of a (*N* −1)×(*N* −1) matrix, which is from matrix (*iδ* **I** − **A**) by crossing out its *k*^*th*^ row and *l*^*th*^ column elements. Denoting by 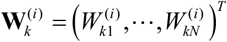 a vector, then the adjacency matrix of (*iδ* **I** − **A**) can be written as

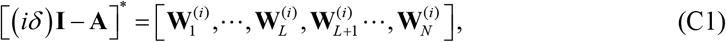

Therefore, it is easy to get the following expression

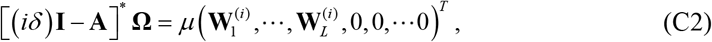

In addition, the determinant is

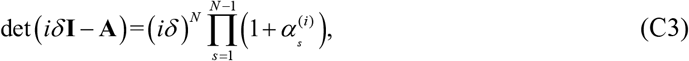

Where 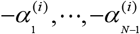 are the nonzero eigenvalues of M-matrix **Ã**^(*i*^) = **A**/*iδ*. Substituting Eqs. (C2), and (C3) into expression 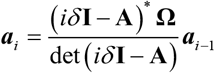, which can be derived from Eq. (9) in the main text, we can obtain the expressions of the factorial BMs

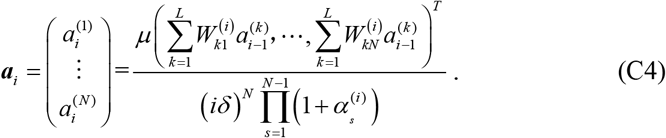

Finally, we can get the expression of the total BMs

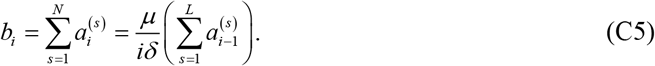

Eqs. (C4) and (C5) imply that the vectors of BMs ***a***_*i*_ and total BMs *b*_*i*_ can be analytically given based on ***a***_0_, i.e., on the steady-state probabilities of promoter states.

### D. Derivation the explicit expression of generating function for the mixture loop model

For the mixture loop model, matrix **H** (*z*) becomes

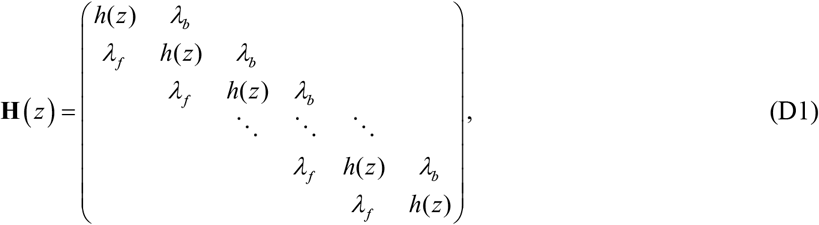

Where *h* (*z*) = *µ* (*z* −1) − *λ*_*f*_ −*λ*_*b*_. Note that matrix **H** (*z*) has eigenvalues 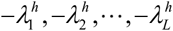 with

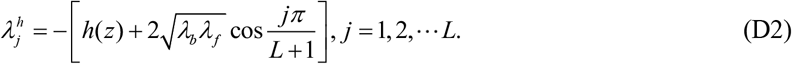

In addition, the eigenvectors associated with the eigenvalues 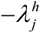 are given by 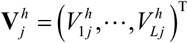 with 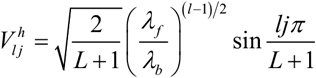. If denoting by 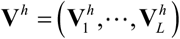, then we have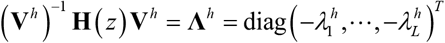. Making the transform 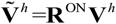, and we obtain the orthogonal matrix 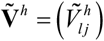 with 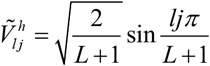. Substituting this expression and Eq. (D2) into Eq. (13) in the main text, we obtain the explicit expression of generating function in the case of the mixture mechanism.

## Notes

### Competing Interest Statement

The authors have declared no competing interest.

